# Loss of zebrafish *dcst2* expression is not associated with muscle abnormalities

**DOI:** 10.1101/2023.08.03.551814

**Authors:** X. Allard-Chamard, E.C. Rodríguez, B. Brais, G.A.B. Armstrong

## Abstract

In this study we examined if the gene encoding Dendritic Cell-specific Six Transmembrane domain containing protein 2 (*dcst2*) plays a role in vertebrate muscle biology. Using the CRISPR/Cas9 mutagenic system we generated a 2 nucleotide deletion in exon 3 of the zebrafish ortholog *dcst2* which resulted in a premature stop codon. Homozygous carriers of the mutation displayed reduced transcriptional expression of *dcst2* suggesting that our mutation was indeed disrupting gene function. Mutant *dcst2* zebrafish developed normally to adulthood and displayed no differences in motor function using a free-swim and swim tunnel assays. Furthermore, histological examination of muscle cells revealed no differences in slow-twitch or fast-twitch muscle cell cross-sectional area in our mutants. We did observe that *dcst2*^-/-^ zebrafish were slightly heavier in weight and males were infertile. The data collected here, suggest that *dcst2* does not play a role in zebrafish muscle cell biology.

## Introduction

Muscle hypertrophy has been documented in several myopathies including non-dystrophic myotonias (Trip et al., 2009), anoctaminopathy (limb girdle muscular dystrophy 2L) (Little et al., 2013), McArdle disease (Quinlivan et al., 2010), Brody disease (Molenaar et al., 2020), and rippling muscle disease (Torbergsen, 2002). Genes associated with these disorders include *CLCN1, SCN4A, ANO5, PYGM, RYR1, SERCA1, CAV3*. and *MSTN* (Bayona et al., 2011; Fermin et al., 2011; Snoeck et al., 2015; Scalco et al., 2016; St-Denis et al., 2016; Walters, 2017; Witting et al., 2018). Recently, using a combination of linkage analysis and exome sequencing, a c.2276T>C missense mutation (encoding the L759P variant) was identified in *DCST2* that segregated with a supranormal strength, muscle hypertrophy and myalgia in a large French-Canadian family (Brais et al., 2016). *DCST2* encodes the protein termed Dendritic Cell-specific Six Transmembrane domain containing protein 2 and is situated on chromosome 1, containing 15 exons. As zebrafish have been previously used to explore muscle disorders (Bassett and Currie, 2003; Guyon et al., 2003; Guyon et al., 2005; Moore et al., 2008; Thornhill et al., 2008; Guyon et al., 2009; Kawahara et al., 2010) we rationalized that generating a zebrafish *dcst2* knockout model may further our understanding of its potential role in muscle biology.

## Materials and Methods

### Zebrafish husbandry and ethical considerations

Wild type (TL) zebrafish (*Danio rerio*) were maintained and bred according to standard procedures (Westerfield, 1995) at the Centre for Neurological Research Disease Models at the Montreal Neurological Institute of McGill University in Montréal, Québec, Canada. Animals were raised in 6 litre tanks at 28.5 °C under a 14/10-hour light/dark cycle. All experiments were performed in accordance with the guidelines of the Canadian Council for Animal Care and approved by the Animal Care Committee of the Montreal Neurological Institute.

### CRISPR mutagenesis

Synthesis of Cas9 mRNA and guide RNA (gRNA) were performed as previously described (Jao et al., 2013; Vejnar et al., 2016). Briefly, a zebrafish codon optimized Cas9 (pT3TS-nCas9n, Addgene plasmid # 46757) was linearized with XbaI overnight and 1 μg of linear template DNA was used for *in vitro* transcription of mRNA using the T3 mMESSAGE mMACHINE® Kit (Invitrogen) followed by phenol-chloroform extraction and precipitation with ethanol. The selected gRNA target site was identified using CRISPRscan (Moreno-Mateos et al., 2015) and synthesized using the T7 MEGAscript kit (Invitrogen) and purified by phenol-chloroform extraction and ethanol precipitation. The following *dcst2* gRNA target site was used (PAM site is underlined): <u>GGT</u>CTGTGTTGTTGAGGACGTCG. A 0.5 nL solution containing *Cas9* mRNA (100 ng/μL), sgRNA (100 ng/μL), and fast green (0.05%, Sigma-Aldrich) was injected into the one-cell stage zebrafish embryo. The primers used to identify zebrafish with indels at the gRNA target loci were AGACAAAGTCCTTCCTCATGTGAC for the forward primer and AAAGGTGCATTGTTGAT GTCATTGAC for the reverse primer and the amplicon was sequenced by Genome Quebec.

### Real-time quantitative PCR

The *dcst2* expression levels were assessed by RT-qPCR. Briefly, complementary DNA (cDNA) derived from RNA from the testis was used as a reference expression control for cDNA obtained from trunk muscle. The primer sequences used were GCTCTCACCTACATCAGGCCC for the forward primer and ACTGGAGCTGCTTGTGCCACGAT for the reverse primer amplifying a product 197 nucleotides in length. For a reference control, *actb2* was used, using the following sequences: CGAGCT GTCTTCCCATCCA for the forward primer; and TCACCAACGTAGCTGTCTTTCT for the reverse primer amplifying a product 86 nucleotides in length. RT-qPCR were performed using SYBR Green (Life Technologies) and run on a LightCycler 96 (Roche).

### Zebrafish motor behaviour

Adult locomotor behaviour was assessed in wild type siblings, heterozygous and homozygous carriers of the mutant allele. Swim distance and speed for each animal was calculated over a 10-minute time period in an aquatic arena (30 cm by 30 cm). Data was collected using a camera (IDS uEye, IDS Imaging Development Systems) and swim behaviour was processed using a tracking system (EthoVision XT, Noldus Information Technology). To further investigate swim behaviour zebrafish were placed in a swim tunnel (Loligo Systems) and maximum swim speed (Umax) was determined using previously established methods for fish (Farrell, 2008). In brief, U_max_ was calculated as the water velocity at which the fish fatigued when the water velocity was increased by 5 cm/minute.

### Tissue collection

Adult zebrafish were euthanized by immersion in 4 °C fish water. The body of the fish was dissected and segmented according to (Benedetti et al., 2016). Muscle tissue, from the trunk region between the anal fin and the base of the caudal fin was selected. Trunks were then mounted in Tragacanth gum (Sigma-Aldrich) and snap frozen in 2-methylbutane (Sigma-Aldrich) chilled in a bath of liquid N_2_ (12-20 seconds).

### Muscle histology characterization using H&E staining

Zebrafish sections of 10 μm thickness were transferred to slides for nuclear (hematoxylin) and cytoplasmic (eosin) staining. Mounted muscle slices were first washed with 1X PBS (137 mM NaCl, 2.7 mM KCl, 10 mM Na_2_HPO_4_, 2 mM KH_2_PO_4_) for 2 minutes followed by fixation using 4% PFA in PBS for 10 minutes. Following this, slides were rinsed with PBS for 2 minutes and then washed with hematoxylin for 1 minute and rinsed with running water. Slides were then treated with a solution of 1% HCl in 10% EtOH for 10 seconds followed by a solution of 80% EtOH (10 dips) and then stained with 0.25% Eosin (1% Eosin in 80% EtOH and Glacial Acetic Acid) for 10 minutes. Finally, the tissue was dehydrated by exposing the slides to an increasing concentration of ethanol for 2 minutes each: 30%, 50%, 70%, 100%, 100%, and then Xylene.

### Sperm morphology and motility

The sperm morphology and motility were assessed at 10X and 40X using a Axio Examiner.A1 microscope (Zeiss). Approximately 0.2 μL of sperm were collected per fish and around 100 μL of system water were added on the slide to activate it. Motor activity was recorded using a camera (Flea3, FLIR) attached to the microscope.

### Statistical analysis

All statistical tests were performed using Prism 9. A Shapiro-Wilk test was performed to assess normality. Statistical differences were determined by *t*-test, one-way ANOVA test and chi square. A *p* value of < 0.05 was considered significant for all statistical testing. Cross-sectional area of muscle fibers was calculated using Fiji.

## Results

To disrupt zebrafish *dcst2* we used the CRISPR/Cas9 system in recently fertilized embryos to generate indels at the target locus and selected a founder with a two base pair deletion in exon 3 (**Fig. 1**). This frameshift mutation resulted in a putative premature stop codon. The founder line was raised and outcrossed and heterozygous offspring were then incrossed as this would permit the generation of homozygous, heterozygous carriers of our mutation as well as wild type zebrafish (**Fig 2A**). However, the genotypic ratio generated from our incross deviated from the predicted 0.25:0.50:0.25 ratio and was 8:54:25 (n = 87) according to homozygous (*dcst2*^-/-^), heterozygous (*dcst2*^+/-^), and wild type (*dcst2*^+/+^) animals respectively, and was significantly different from Mendelian proportions (χ2 = 11.71, n = 87, *p* < 0.01) (**Fig. 2B**).

**Fig 1.**
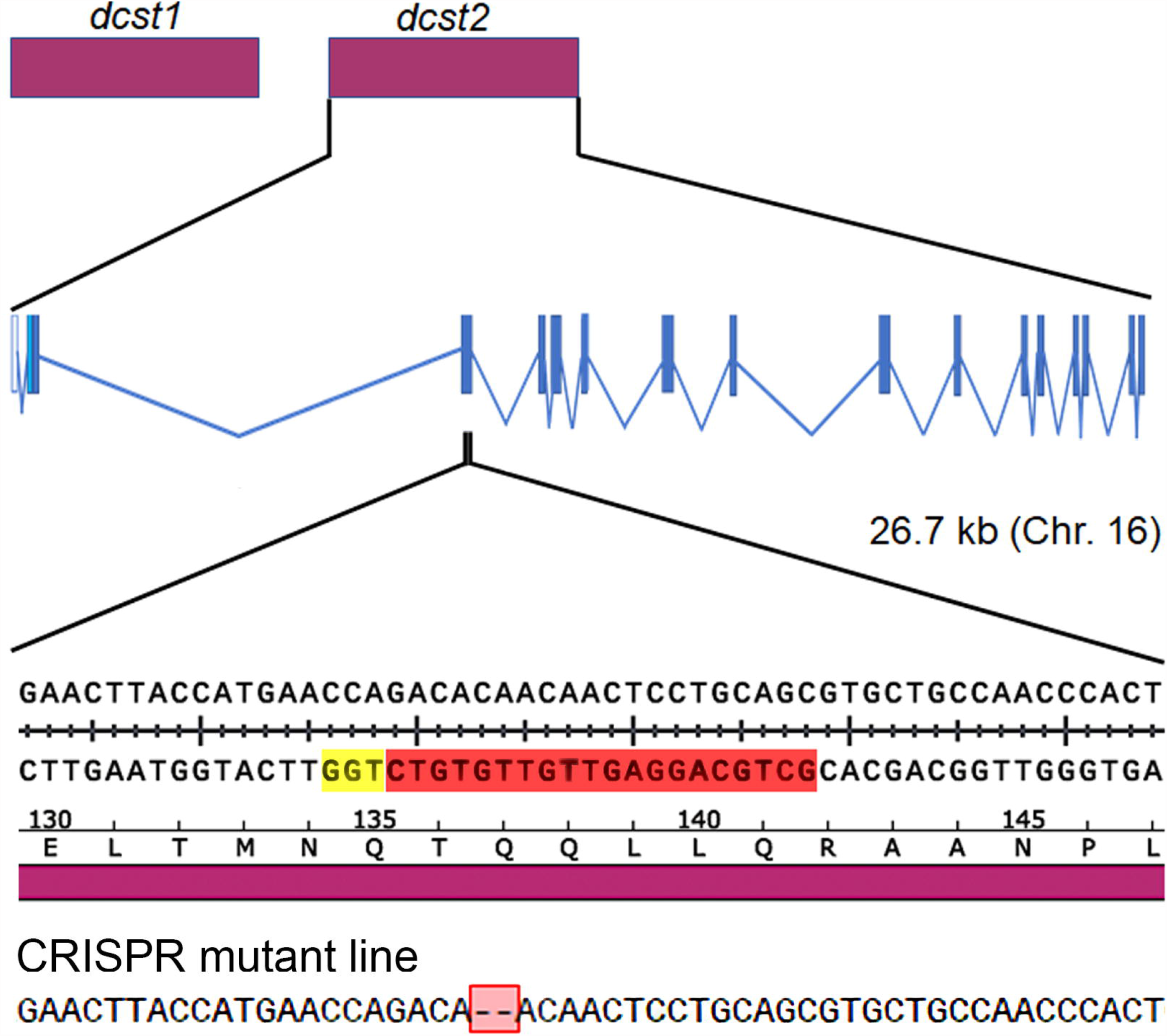
Genomic structure of *dcst1* and *dcst2* in the zebrafish genome and the CRISPR gRNA target site used to generate a knockout mutant. Zebrafish *dcst2* contains 16 exons and is located on chromosome 16. Blue exons denote translated exons and white exons are the Untranslated Region (UTR). A frame shift mutation in exon 3 was identified in a founder mutant line resulting in a premature stop codon. Yellow nucleotides indicate PAM sequence and red nucleotides indicate the guide RNA target sequence.

**Fig 2.**
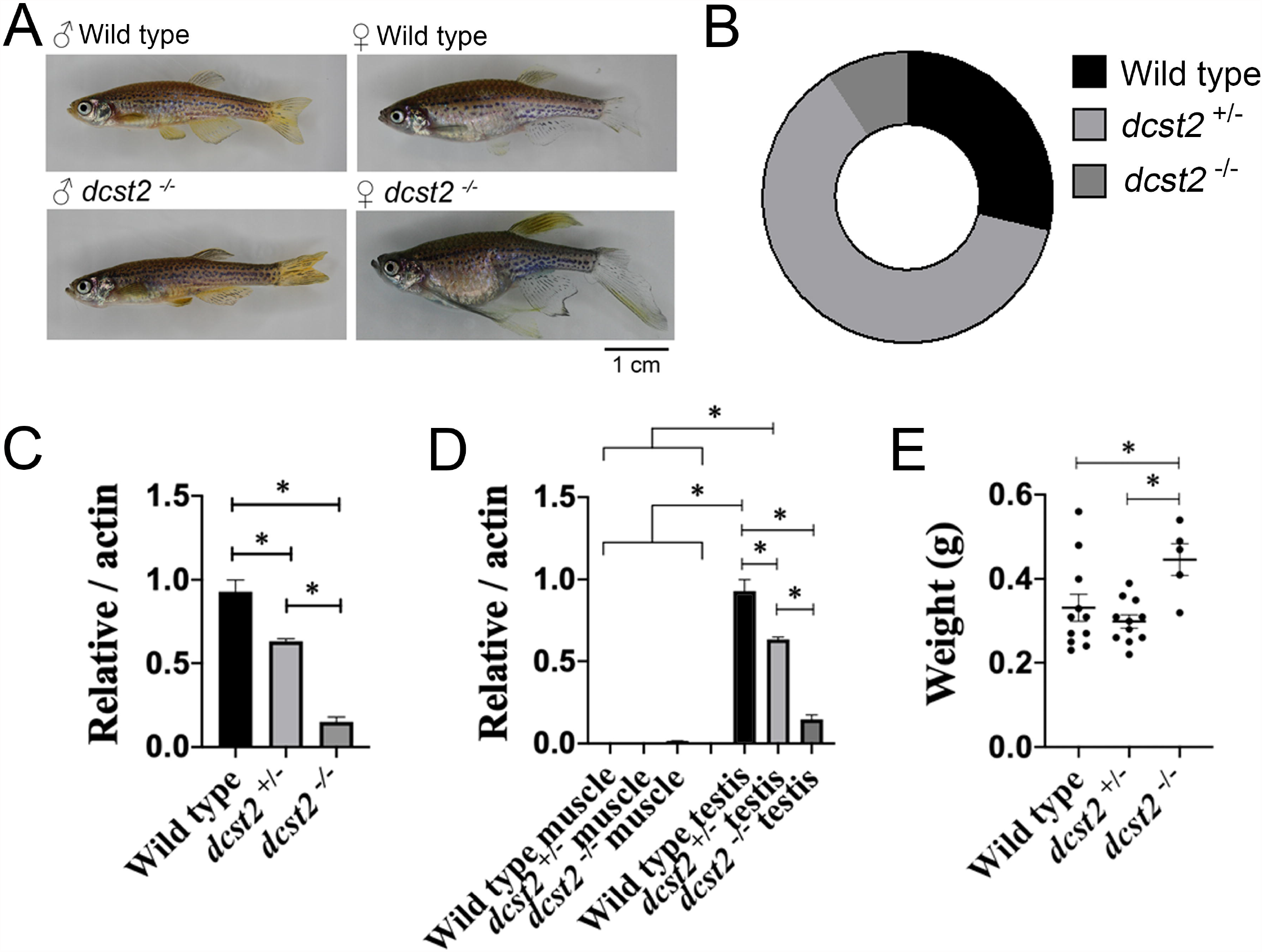
Example images of male and female adult zebrafish aged 12 months (**A**). Scale bar represents 1 cm. Genotypic ratios generated from crosses between two *dcst2*^+/-^ mutant zebrafish depicted as a pie chart (**B**). Transcriptional expression levels of *dcst2* relative to *actin* in the testis confirming that our mutation likely is being degraded by nonsense-mediated decay (**C**). Asterisks represent a statistically significant difference (*p* < 0.05). **D**, A comparison between muscle and testis *dcst2* expression. Asterisks represent a statistically significant difference (*p* < 0.05). **E**, Tabulation of male body weight of each fish from each genotype. Asterisks represent a statistically significant difference (*p* < 0.05).

While our main interest was to examine if disruption of *dcst2* conferred a muscle cell phenotype, a recent study suggested that it plays a role in acrosomal fusion of sperm with oocytes (Noda et al., 2022). To examine if a similar phenomenon was occurring in our fish models we tested if male *dcst2*^-/-^ could fertilize wild type zebrafish eggs. Mutant *dcst2*^-/-^ male zebrafish could reliably flush eggs from a wild type female but none of the eggs were fertilized. Conversely, female *dcst2*^-/-^ eggs could be fertilized using wild type males. We then examined if there was a sperm motility defect in our *dcst2*^-/-^ males. Examination of sperm from all three genetic groups did not reveal any obvious changes in sperm motility suggesting that the fertility defect is not associated with poor sperm motor function. Our data can be well explained by the research findings recently published by Noda and colleagues confirming that Dcst2 is involved with acrosomal fusion with oocytes (Noda et al., 2022).

As our main focus was on investigating the impact of loss of *dcst2*^-/-^ on muscle function, we next wanted to confirm that our frameshift mutation resulted in reduced expression of the *dcst2*^-/-^ transcript. We used cDNA derived from RNA collected from the testis to measure the levels of the *dcst2* transcript and found that cDNA from mutants was reduced (one-way ANOVA test, F (2, 5) = 123.5, *p* < 0.05) suggesting that the transcript was sent for nonsense mediated decay (**Fig. 2C**). The expression of *dcst2* in trunk muscle was not high enough to distinguish variation across our genetic groupings and testis expressed significantly much more *dcst2* than muscle (one-way ANOVA test, F (5, 8) = 178.2, *p* < 0.05) (**Fig. D**). Examination of the gross morphology of our genetic groupings revealed that the weight of *dcst2*^-/-^ adult zebrafish was significantly higher than *dcst2*^+/-^ and wild type zebrafish (one-way ANOVA test, F (2, 24) = 123.5, *p* < 0.05) (**Fig 2E**).

We next examined if motor function was altered in our mutant models. Adult zebrafish were placed in an aquatic arena and motor activity was recorded for 5 minutes. No significant differences in swim distance or mean swim velocity were observed across our genetic groupings (**Fig 3A-C**). Similarly, measurement of the swimming speed (U_max_) using a swim tunnel did not reveal any differences in motor function in our mutant *dcst2* model (**Fig 3 D, E**). While we did not observe any defects in motor function in our mutant *dcst2* model we nevertheless examined if there were any abnormalities in the trunk musculature. However, we did not observe muscle fiber hypertrophy, nor did we observe muscle atrophy in our *dcst2*^-/-^ model. Specifically, examination of slow-twitch and fast-twitch muscle fiber cross-sectional areas revealed no significant differences between our *dcst2*^*-/-*^ and wild type muscle cells (**Fig 4A-D**).

**Fig 3.**
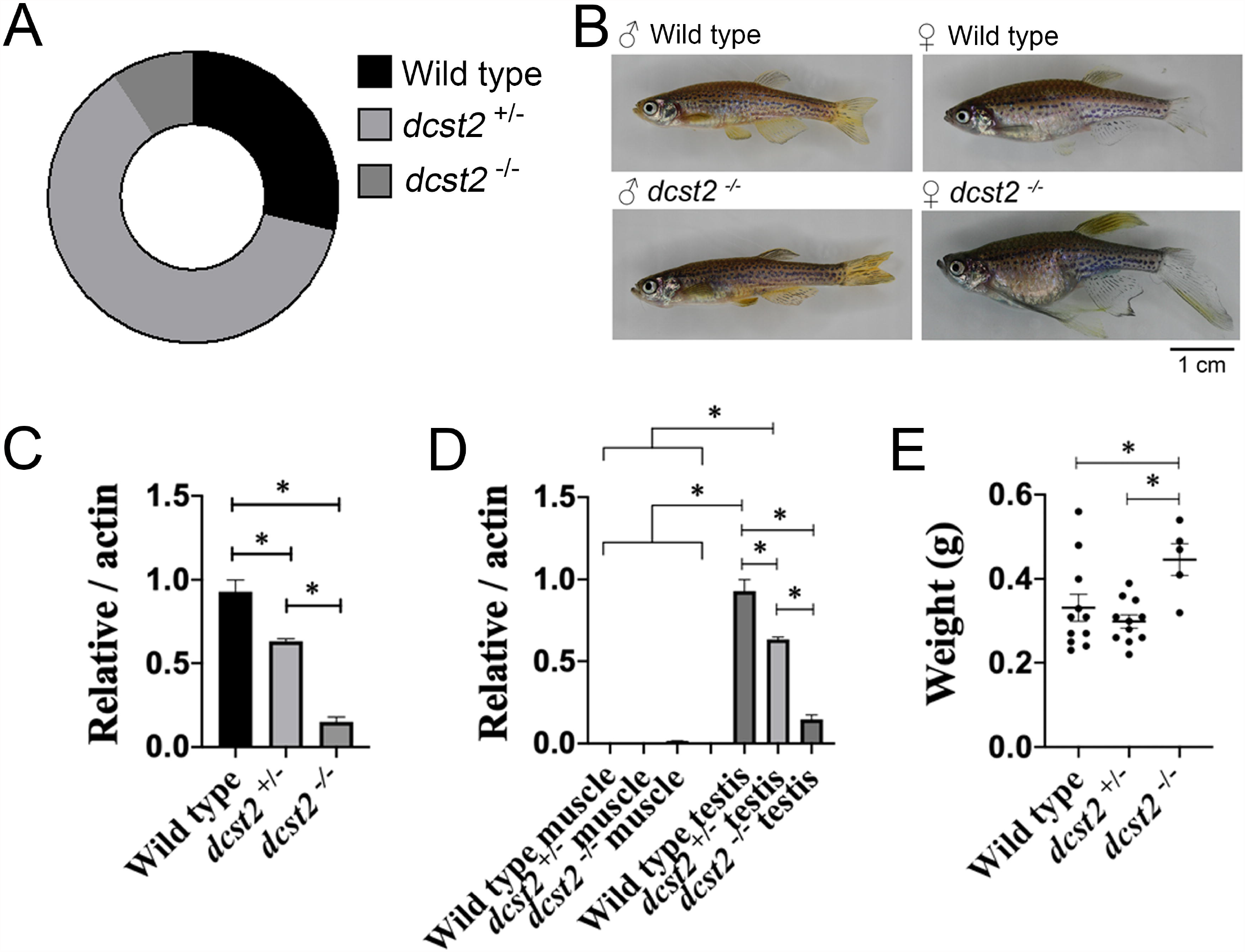
Example heat maps of individual fish swimming behaviour in an open field aquatic arena (**A**). Swim distance (**B**). Mean swim velocity of open field swimming behaviour (**C**). Example image of the swimming tunnel used to assess muscle performance of fish (**D**). Tabulation of the maximum swim speed (U_max_) of our genetic groupings (E). No significant differences were found in any of our measures of motor function.

**Fig 4.**
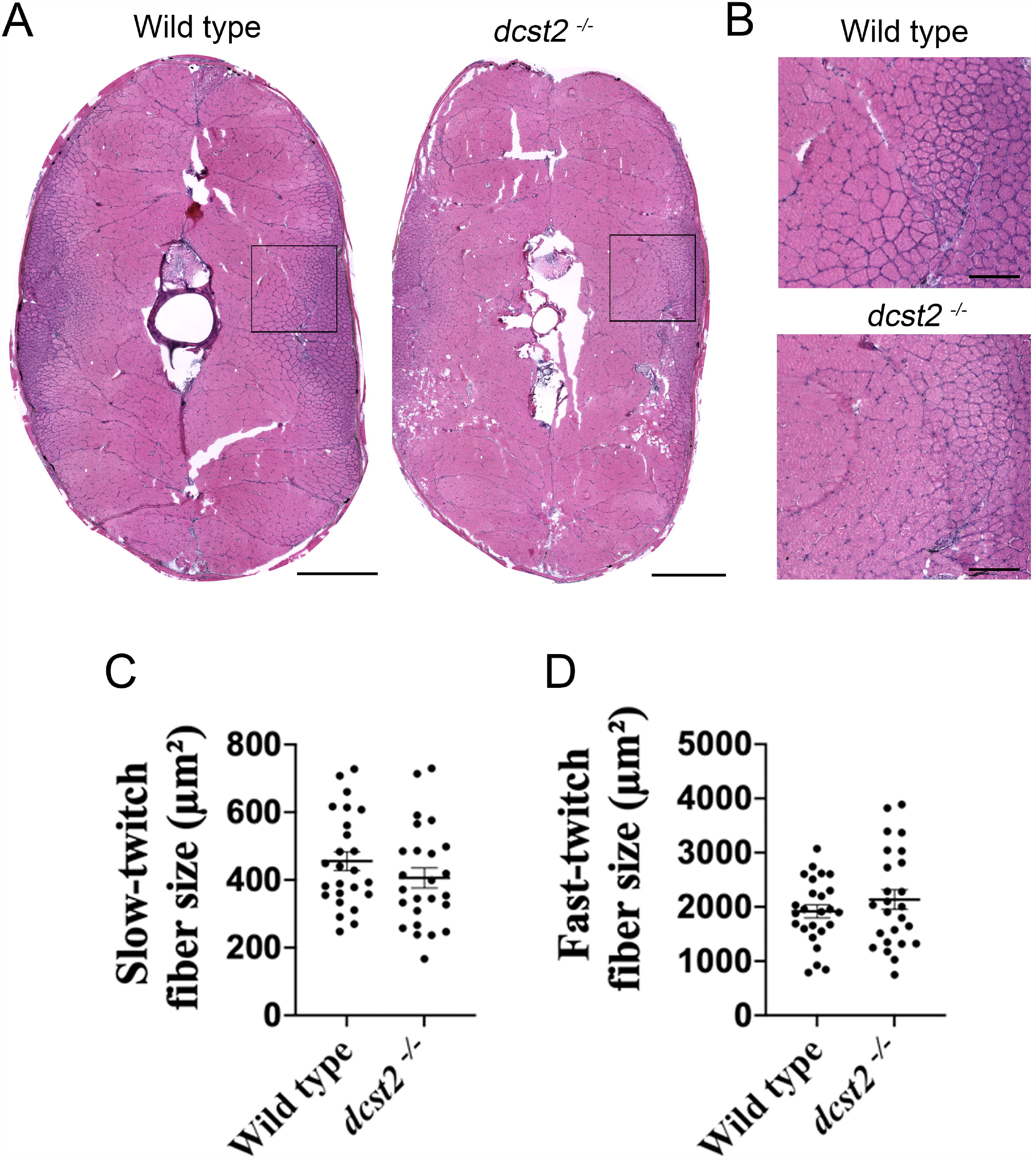
Hematoxylin and eosin staining of zebrafish trunk cryosections (**A**). Scale bar represents 500 μm. Insets of cross sections showing no obvious muscle fiber abnormalities (**B**), scale bar represents 50 μM. Tabulation of slow-twitch muscle fiber cross-sectional area between wild type and *dcst2*^-/-^ muscle cells (**C**). Tabulation of fast-twitch muscle fibers cross-sectional area wild type and *dcst2*^-/-^ zebrafish (D). No significant differences were found in any of our measures of muscle cells.

## Discussion

While we originally had speculated that *DCST2* may be involved with muscle hypertrophy and strength, the data presented here does not support that hypothesis. Using the CRISPR/Cas9 mutagenic system we generated a zebrafish *dcst2*^-/-^ knockout model. We observed no differences in motor function in adult *dcst2*^-/-^ zebrafish when compared to wild type siblings. Moreover, the absent expression of *dcst2* in trunk musculature and lack of muscle fiber hypertrophy in *dcst2*^*-/-*^ zebrafish further suggests that this gene is unlikely to play a significant role in muscle biology. We did observe two findings of note. The first being that *dcst2*^-/-^ zebrafish were slightly heavier than heterozygous KO and wild type siblings (**Fig 2E**). A genome-wide association study did identify a rare single nucleotide polymorphism in *DCST2* that is associated with height in humans (van der Valk et al., 2015). *DCST2* is an important regulator of osteoclast cell fusion in bone homeostasis (Kukita et al., 2004; Yagi et al., 2005) and it may be that variants alter osteogenesis during growth. We speculate that loss of *dcst2* may confer a slight but significant change in the size of zebrafish. The second observation made by our group was that sperm from male *dcst2*^-/-^ fish were unable to fertilize eggs. This finding is in keeping with previous data showing that both Dcst1 and Dcst2 are essential for the binding of sperm to the oolemma (Noda et al., 2022). Although our main interest was to explore *DCST2* potential in muscle biology and function, our findings suggest that it likely does not play a significant role in this tissue.

